# Deazaguanylation is a nucleobase-protein conjugation required for type IV CBASS immunity

**DOI:** 10.1101/2025.04.06.647259

**Authors:** Douglas R. Wassarman, Patrick Pfaff, Joao A. Paulo, Steven P. Gygi, Kevan M. Shokat, Philip J. Kranzusch

**Affiliations:** Department of Microbiology, Blavatnik Institute, Harvard Medical School, Boston, MA, USA; Department of Cancer Immunology and Virology, Dana-Farber Cancer Institute, Boston, MA, USA; Department of Cellular and Molecular Pharmacology and Howard Hughes Medical Institute, University of California, San Francisco, CA, USA; Department of Cell Biology, Blavatnik Institute, Harvard Medical School, Boston, MA, USA; Parker Institute for Cancer Immunotherapy at Dana-Farber Cancer Institute, Boston, MA, USA

## Abstract

7-deazapurines are nucleobase analogs essential for nucleic acid modifications in nearly all cellular life. Here, we discover a role for 7-deazapurines in protein modification within type IV CBASS anti-phage defense and define functions for CBASS ancillary proteins Cap9 and Cap10 in nucleobase-protein conjugation. A structure of Cap10 reveals a tRNA transglycosylase-family enzyme remodeled to bind the modified N-terminus of a partner cGAS/DncV-like nucleotidyltransferase linked to a 7-amido-7-deazaguanine (NDG) nucleobase. The structure of Cap9 explains how this QueC-like enzyme co-opts a 7-deazapurine biosynthetic reaction mechanism for NDG conjugation. We demonstrate that Cap9, Cap10, and NDG conjugation are essential for host defense against phage infection. Our results define a previously unknown 7-deazapurine protein modification and explain how nucleobase biosynthetic machinery has been repurposed for antiviral immunity.

## MAIN TEXT

7-deazapurines are nucleobase analogs that control essential cell processes in all domains of life (*1*). Carbon substitution at the N7 nitrogen of purine bases creates a chemical scaffold that is readily modified into distinct 7-deazapurine compounds with broad roles in RNA and DNA modification (*2*–*4*), in natural antibiotics including toyocamycin and sangivamycin (*5*), and in other secondary metabolites (*6, 7*). First discovered in 1972, a 7-deazaguanine nucleobase named Q (queuine) is post-transcriptionally incorporated into the anticodon loops of tRNAs in bacterial, plant, and animal cells (*2, 8*). Q incorporation directly alters codon preference and is essential for maintaining cellular translation efficiency (*5*–*11*). Q precursors can additionally be incorporated into bacterial DNA (*3*) and are used by phages to evade bacterial immunity (*4*).

In bacteria, biosynthesis of Q requires a set of dedicated proteins that includes the enzyme QueC (7-cyano-7-deazaguanine synthase) responsible for generating the intermediate nucleobase preQ_0_ (*12, 13*). Q, preQ_0_, and other 7-deazaguanines are installed into tRNA and nucleic acid targets by the protein TGT (tRNA-guanine transglycosylase) and related transglycosylase enzymes (*3, 14, 15*). Eukaryotes acquire Q from diet and gut microbiota (*16*) and install Q into tRNA through the activity of genomically encoded eukaryotic TGT homologs. Notably, Q biosynthetic genes have been recently identified in various bacterial anti-phage defense systems (*17*–*19*) suggesting potential unexplained roles for 7-deazaguanines in antiviral immunity.

### QueC and TGT homologs are required for bacterial defense against diverse phages

Q biosynthetic enzymes are frequent components of anti-phage defense systems named CBASS (cyclic oligonucleotide-based antiviral signaling systems) that synthesize cyclic dinucleotide and cyclic trinucleotide immune signals in response to phage infection (*17, 19*). All CBASS operons encode a CD-NTase (cGAS/DncV-like nucleotidyltransferase) enzyme that catalyzes nucleotide immune signal synthesis in response to infection and a Cap (CD-NTase-associated protein) effector that restricts phage replication (*20*–*22*). Type IV CBASS operons additionally encode accessory proteins Cap9 and Cap10 that are predicted to share homology with Q biosynthetic enzymes QueC and TGT (*17, 19*). Approximately one-third of type IV systems also encode Cap11, a protein related to the DNA glycosylase OGG (Fig. 1A). Type IV CBASS operons occur in defense islands across diverse bacterial phyla and archaea, suggesting a broadly conserved role in antiviral immunity (Fig. 1A, 1B).

**Figure 1.**
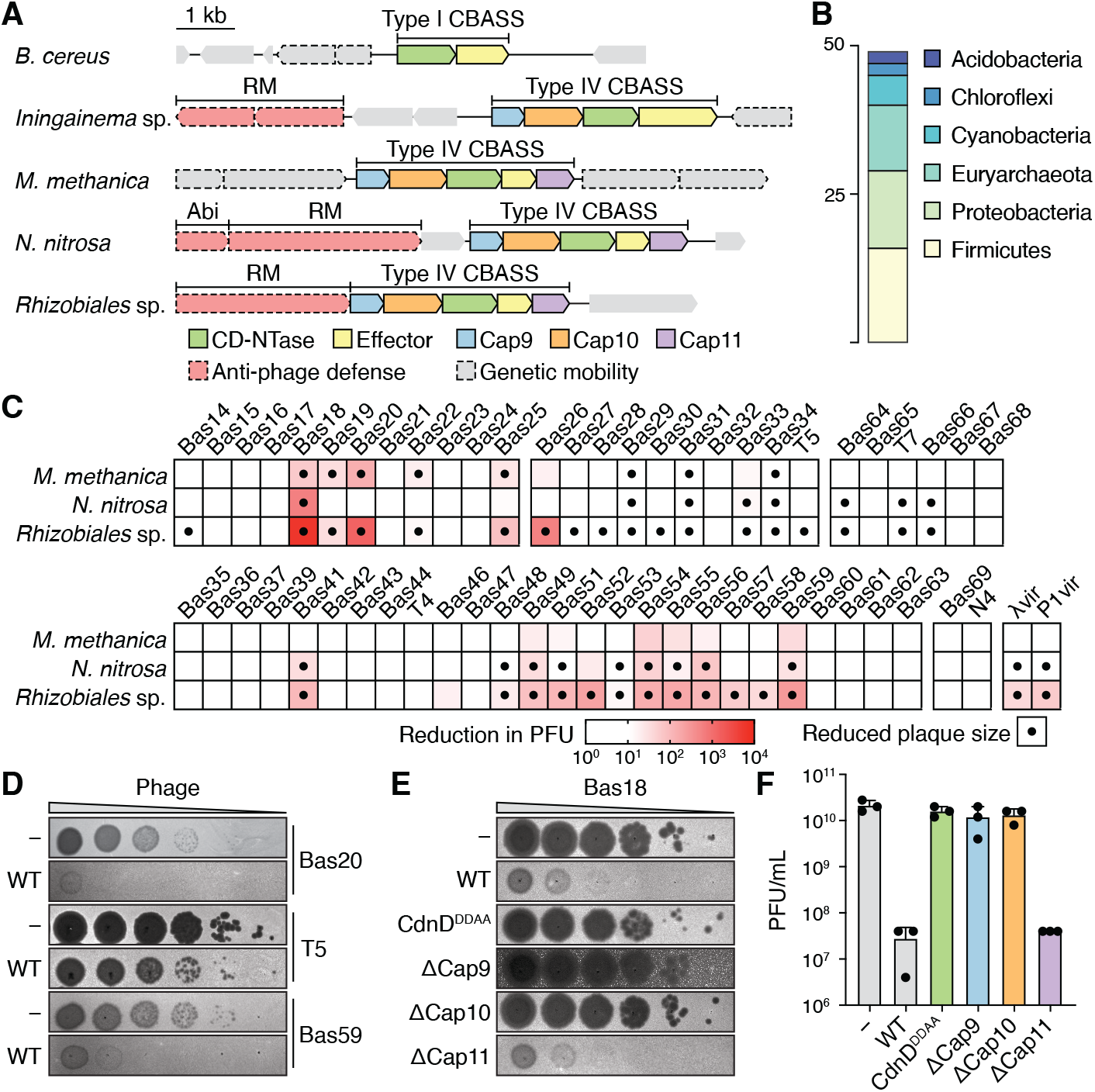
Type IV CBASS defends bacteria against diverse phages. (**A**) Genomic context of representative CBASS operons. See Table S1 for genome accession numbers. (**B**) Distribution of host organism phyla for 49 identified type IV CBASS operons. See Table S1 for genome accession numbers. (**C**) Anti-phage activity of type IV CBASS operons expressed in *E. coli* induced with 0.2% arabinose. Data represent the reduction in plaque forming units (PFU) or plaque size compared to bacteria expressing a GFP control vector (n=2). Phages are grouped by family. (**D**) Representative plaque assays with *E. coli* expressing a GFP control (−) or wildtype *Rhizobiales* CBASS operon (WT) induced with 0.2% arabinose and challenged with the indicated phages. (**E**) Representative plaque assays with *E. coli* expressing GFP control (−) or *Rhizobiales* CBASS operons with the indicated genotypes and challenged with phage Bas18. Wildtype (WT), CdnD D94A/D96A (CdnD^DDAA^). (**F**) Quantified plaque forming units (PFU) from plaque assay as in (E), n=3.

To test the defensive capability of type IV CBASS, we expressed operons from three proteobacterial species in *E. coli* and challenged with a diverse panel of 58 phages from six viral families (*23*). All three systems inhibited phage infection, including up to 5,000-fold reduction for phages Bas18 and Bas20 and diminished plaque size for many phages including T5 and T7 (Fig. 1C). We selected a *Rhizobiales* operon exhibiting robust defense for mechanistic characterization (Fig. 1C, 1D). Mutation of conserved metal-coordinating residues in the *Rhizobiales* CD-NTase enzyme CdnD (D94A/D96A), or deletion of either Cap9 or Cap10, completely abolished anti-phage defense, while deletion of Cap11 had no effect (Fig. 1E, 1F). Deletion of Cap9 or Cap11 increased cellular toxicity in the absence of infection, further demonstrating a key role of these accessory proteins in controlling immune activation (Fig. 1E, Fig. S1). These results establish type IV CBASS as a widespread anti-phage defense system that requires both Cap9 and Cap10 proteins to protect against viral infection.

### Cap10 forms an essential complex with CdnD through remodeled interfaces

To define the role of Q biosynthetic proteins in anti-phage defense, we purified individual protein components from cells expressing type IV CBASS. Immunoprecipitated CdnD co-purified with a second protein corresponding to the predicted molecular weight of Cap10 (Fig. 2A). We determined a 2.4 Å X-ray crystal structure of the purified complex revealing a symmetric 2:2 assembly with a core homodimer of two Cap10 proteins flanked by two CdnD protomers (Fig. 2B, Table S2). The Cap10 structure adopts a canonical TGT-like transglycosylase fold composed of a TIM-like (αβ)_8_-barrel (*15*). Cap10 exhibits high structural similarity to the human enzyme QTRT1, confirming clear homology with Q biosynthesis machinery (Fig. S2) (*24*). The structure of CdnD exhibits the characteristic nucleotidyltransferase fold of other CD-NTases, with the active site formed between an α/β N-terminal lobe and C-terminal α-helix bundle (Fig. S3) (*20*). However, each molecule of CdnD in complex with Cap10 adopts a conformation distinct from previously observed CD-NTase structures. Superposition of the *Rhizobiales* CdnD structure from type IV CBASS with an *Enterobacter cloacae* CdnD structure from a type II CBASS operon (*25*) reveals a 30° rotation and 7.5 Å displacement of the N-terminal lobe (Fig. 2C) that induces bunching at the base of strand β2 and misalignment of conserved metal-binding residues D94 and D96 (Fig. 2D, 2E). Previous studies in *Vibrio cholerae* identified a regulatory site on the CD-NTase surface opposite the active site where folate metabolites bind and inhibit enzyme activity (*26*–*28*). In the Cap10–CdnD structure, we observed coenzyme A bound in an adjacent site on each CdnD protomer, further suggesting that CdnD is in an inactive conformation (Fig. S3). The coenzyme A forms a disulfide bridge with CdnD residue C191, but this interaction appears nonessential for function, as C191 is poorly conserved and C191A mutation had no impact on type IV CBASS anti-phage defense (Fig. S3). Consistent with these structural observations, we confirmed that the purified Cap10–CdnD assembly does not catalyze cyclic nucleotide synthesis *in vitro* (Fig. S4), supporting that CdnD remains inactive within this protein complex.

**Figure 2.**
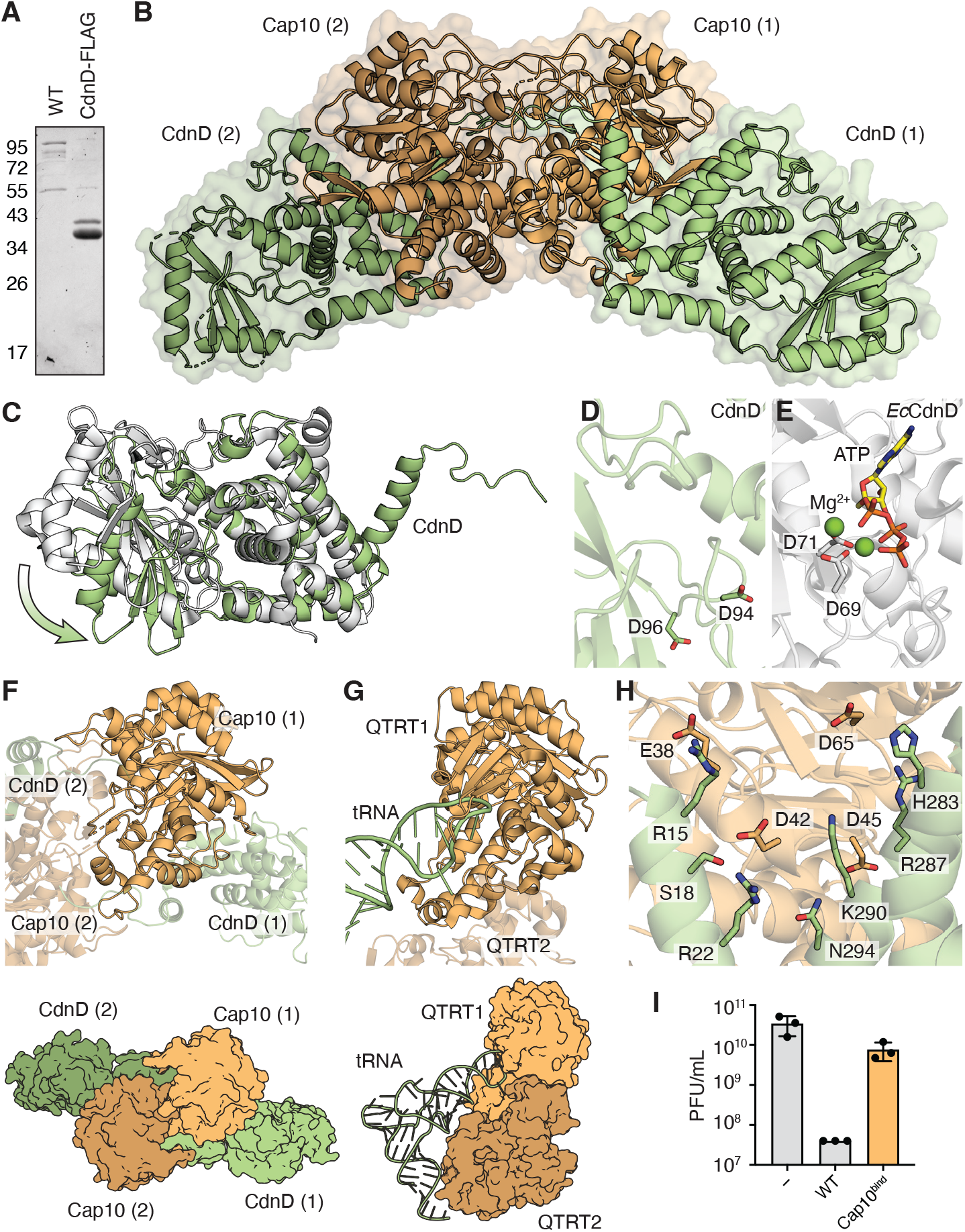
Cap10 is a TGT-family enzyme adapted for protein targeting. (**A**) Coomassie-stained SDS-PAGE following anti-FLAG immunoprecipitation. Lysates were prepared from *E. coli* expressing wildtype (WT) *Rhizobiales* operon or mutant operon encoding a C-terminal FLAG-tagged CdnD (CdnD-FLAG). Calculated molecular weights: CdnD-FLAG (34.5 kDa), Cap9 (19.9 kDa), Cap10 (37.4 kDa), Cap11 (23.5 kDa). (**B**) 2.4 Å X-ray crystal structure of *Rhizobiales* Cap10–CdnD complex. Two protomers of CdnD (green) and two protomers of Cap10 (gold) form a 2:2 heterotetrameric complex. Each Cap10–CdnD dimer is denoted with (1) or (2). See Table S2 for crystallographic statistics. (**C**) Superposition of Cap10-bound CdnD (green) and *Enterobacter cloacae* CdnD (white) aligned by C-terminal lobe shows displacement of N-terminal lobe (arrow). PDB: 7LJL. (**D**) CdnD active site with metal-binding residues D94 and D96. (**E**) *Enterobacter cloacae* CdnD active site with metal-binding residues D69 and D71 (white), Mg^2+^ ions (green), and bound ATP substrate (yellow). PDB: 7LJL. (**F**) and (**G**) Structural comparison of the Cap10– CdnD complex (F) aligned with human QTRT1 in complex with QTRT2 and tRNA (G) demonstrates that canonical nucleic acid interacting surfaces in tRNA transglycosylase-family proteins are remodeled in Cap10 to enable dimerization and CdnD binding. PDB: 8OMR. (**H**) Hydrophilic interface between Cap10 (E38, D42, D45, D65) and CdnD (R15, S18, R22, H283, R287, K290, N294). (**I**) Phage Bas18 plaque forming units (PFU) determined by plaque assay with *E. coli* expressing a GFP control (−) or *Rhizobiales* CBASS operon with the indicated genotypes. Wildtype (WT), Cap10 E38R/D42R/D45R/D65R (Cap10^bind^).

Analysis of the Cap10–CdnD complex reveals extensive structural remodeling that enables Cap10 to dimerize and engage CdnD (Fig. 2F, 2G). The human TGT QTRT1 forms a pseudo-homodimer with its noncatalytic paralog QTRT2 that cooperatively stabilizes tRNA substrate binding in the QTRT1 active site (Fig. 2G) (*24*). In contrast, the tRNA-binding surface is repurposed in Cap10 to create a new Cap10–Cap10 homodimerization interface (Fig. 2F). At this interface, three antiparallel β-strands β7–β9 that normally form a lid over the tRNA substrate are remodeled into loops that interlock between Cap10 protomers (Fig. 2F, 2G, Fig. S2). Superposition of Cap10 and QTRT1 demonstrates clear steric clashes between the partner Cap10 protomer and the canonical nucleic acid binding site that preclude tRNA binding. Cap10 further lacks the Zn^2+^ binding site formed by helices α13–α15 and a positively charged patch near helix α1 that typically stabilize QTRT1-nucleic acid interactions (Fig. S2). The Cap10-CdnD interaction occurs on the opposite protein face at a site that is also distinct from the canonical TGT dimerization interface. Two basic CdnD helices α1 and α13 form a V-shaped clamp that locks around the acidic α1 helix of Cap10 (Fig. 2H). Charge-reversal mutations near this Cap10 helix α1 (E38R/D42R/D45R/D65R) abolish anti-phage defense (Fig. 2I), confirming that Cap10–CdnD complex formation is essential for type IV CBASS activity.

### CdnD is conjugated to a novel 7-deazaguanine modification

The electron density in the Cap10–CdnD complex revealed a surprising conjugation between the N-terminus of CdnD and a modified nucleobase. In the Cap10–CdnD assembly, the N-terminus of each CdnD protomer traverses the complex and binds deep within the opposing Cap10 subunit active site (Fig. 3A). The initial CdnD methionine residue M1 is absent as expected due to normal post-translation processing of bacterial proteins (*29*) and the electron density of the peptide chain begins with glycine residue G2. An F_O_−F_C_ polder omit map reveals clear connective density extending from G2 into a fused planar molecule consistent with a purine nucleobase (Fig. 3B). The CdnD N-terminus is positioned within the Cap10 active site analogous to where TGT enzymes bind Q and related nucleobase precursors, suggesting that the CdnD purine modification is a 7-deazaguanine nucleobase derivative. Of the known 7-deazaguanine derivatives (*1*), the only molecule compatible with the electron density is the nucleobase 7-amido-7-deazaguanine. Notably, highly conserved Cap10 residues D73, S74, D122, and K154 read out each position of the nucleobase through hydrogen bonding interactions, further supporting assignment of this conjugation as 7-amido-7-deazaguanine (Fig. 3B). To verify the CdnD protein modification, we analyzed the nucleobase–protein complex with a combination of intact liquid chromatography-mass spectrometry (LC-MS) and peptide fragmentation LC-MS/MS. Analysis of the intact Cap10– CdnD complex, purified under reducing conditions to remove the coenzyme A molecule, revealed two predominant species: unmodified Cap10 (37,398 Da) and CdnD conjugated to an adduct with a molecular weight of 176 Da (34,576 Da) matching the mass of 7-amido-7-deazaguanine (176.04 Da) (Fig. 3C). Fragmentation analysis using high-resolution LC-MS/MS of trypsin-digested CdnD further confirmed the presence of 7-amido-7-deazaguanine modification on residue G2 (Fig. 3D, Fig. S5), and we named this protein deazaguanylation modification NDG (<u>N</u>-terminal 7-amido-7-<u>d</u>eaza<u>g</u>uanine) (Fig. 3E).

**Figure 3.**
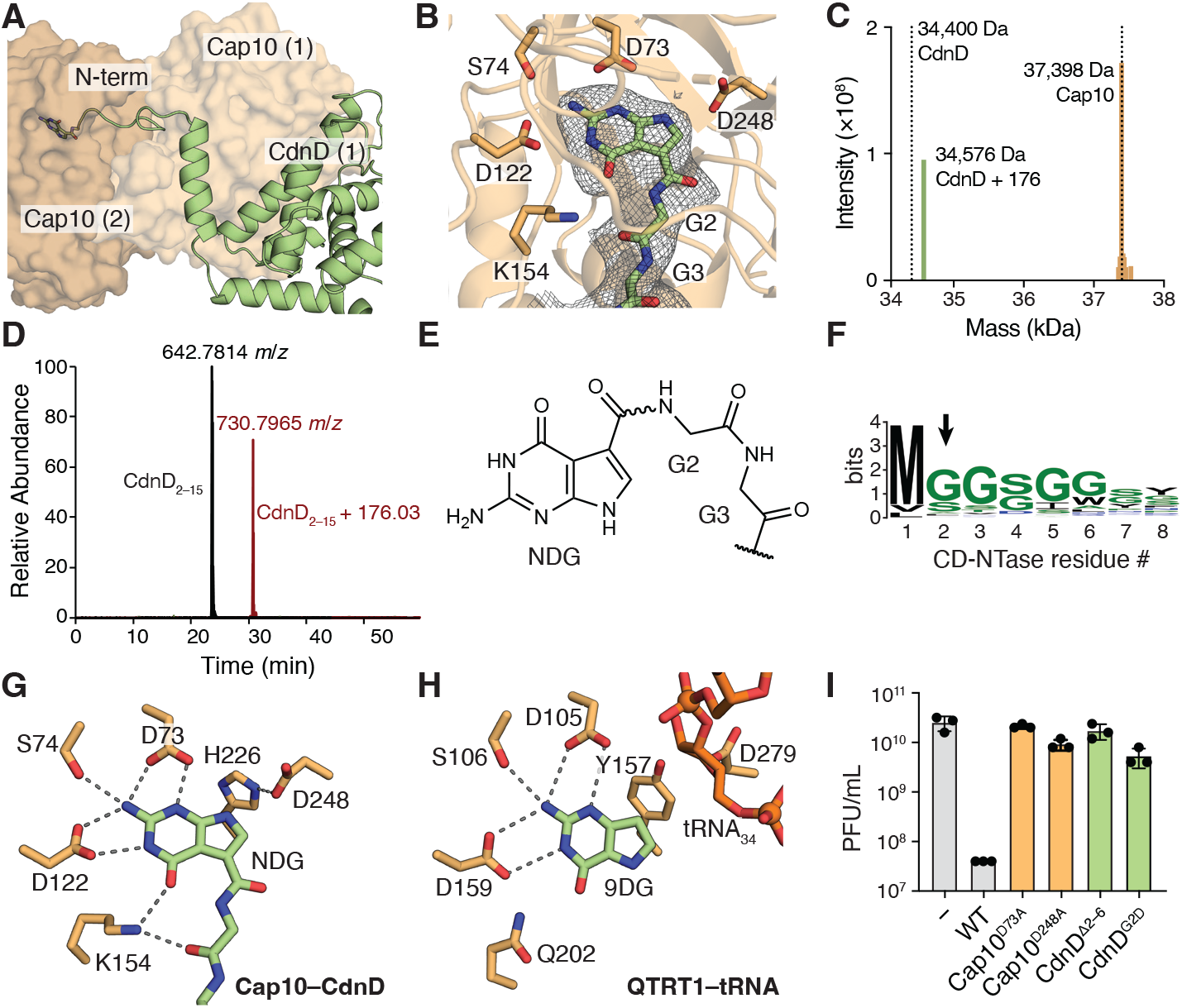
The CD-NTase N-terminus is conjugated to a 7-deazaguanine nucleobase. (**A**) CdnD N-terminal tail (green cartoon) shown binding with the opposite Cap10 protomer (gold surface). CdnD (2) subunit makes a symmetrical interaction with Cap10 (1) but is omitted for clarity. (**B**) CdnD N-terminus with covalent 7-amido-7-deazaguanine nucleobase modification (green) and surrounding Cap10 residues (gold). Gray mesh depicts F_O_−F_C_ polder omit map (sigma=3). (**C**) Intensity of ion masses determined by intact LC-MS analysis and spectral deconvolution of Cap10–CdnD complex. Dotted vertical lines show expected masses of Cap10 and methionine M1-cleaved CdnD. (**D**) Chromatogram of unmodified (black) and modified (red) N-terminal peptides (residues 2–15, GGSGGSFGSGFDPR) from trypsin-digested CdnD. Mass-to-charge ratio (*m*/*z*) of peptide ions with charge 2+ is reported. (**E**) Chemical structure of NDG showing conjugation of 7-amido-7-deazaguanine to glycine 2 of CdnD. Bond connecting NDG and protein N-terminus indicated by wavy line. (**F**) Sequence logo showing conservation of type IV CBASS CD-NTase N-terminus. Black arrow indicates the position of NDG conjugation. (**G**) Cap10 active site showing conserved nucleobase-coordinating residues and catalytic D248/H226 residue pair (gold) and NDG-modified CdnD (green). (**H**) QTRT1 active site showing conserved nucleobase-coordinating residues and catalytic D279/Y157 residue pair (gold), substrate mimic 9-deazaguanine (9DG, green), and trapped tRNA intermediate (orange). PDB: 8OMR. (**I**) Phage Bas18 plaque forming units (PFU) determined by plaque assay with *E. coli* expressing a GFP control (−) or *Rhizobiales* CBASS operon with the indicated genotypes.

Structural analysis of the Cap10–CdnD complex suggests conjugated NDG is poised for transfer onto a target molecule. Comparison of the Cap10 active site with previous substrate-bound structures of a bacterial TGT enzyme (*30, 31*) and human QTRT1 (*24*) reveals a near perfect superposition of NDG and other TGT nucleobase substrates (Fig. 3G, 3H). In QTRT1, the catalytic residue D279, coordinated by hydrogen bonding with nearby Y157, excises guanine from tRNA position 34 through formation of a covalent intermediate with the ribose anomeric carbon, which is resolved by nucleophilic attack from the N9 position of the 7-deazaguanine substrate to generate the modified tRNA product (Fig. 3H) (*31*). The Cap10–CdnD structure shows conserved geometry of an analogous catalytic residue D248 and hydrogen bond donor H226, with D248 poised to transfer the free N9 position of CdnD-linked NDG (Fig. 3G). Furthermore, conserved Cap10 residues D73, S74, D122, and K154 read out the Watson-Crick edge of the NDG nucleobase face, making contacts identical to those made by TGT to recognize 7-deaguanine substrates.

Notably, covalent attachment of a CD-NTase to a target molecule is an essential step controlling activation in type II CBASS anti-phage defense (*32*–*35*). In type II CBASS, the accessory protein Cap2 is structurally homologous to E1/E2-like enzymes in eukaryotic ubiquitin cascades. Cap2 performs a ubiquitin-transfer-like reaction to conjugate AMP to the CD-NTase C-terminus, creating a chemically activated site that can be further transferred to a target molecule upon phage infection. Cap10-mediated transfer of NDG represents a chemically distinct but functionally similar process, potentially enabling CD-NTase attachment to target molecules in type IV CBASS defense. Supporting this hypothesis, we mutated the Cap10 nucleophilic residue D248, or nucleobase recognition residue D73, and observed complete loss of anti-phage defense (Fig. 3I). Furthermore, we found that each feature of NDG conjugation is conserved throughout type IV CBASS operons, including universal presence of a CD-NTase N-terminal extension that begins with the M(G/S)G(G/S)GG motif where NDG conjugation occurs (Fig. 3F, Fig. S6). Deletion of this motif in *Rhizobiales* CdnD, or substitution of residue G2 with a bulky aspartic acid mutation, abolished all anti-phage defense (Fig. 3I). Despite significant effort from several groups, the target identity and mechanism of CD-NTase attachment in type II CBASS remains unknown (*32*–*35*). Similarly, we have been unable to identify a target molecule for CD-NTase attachment in type IV CBASS and future experiments will be required to further understand the mechanism of CD-NTase activation. Together our results establish NDG as a novel post-translational modification required for type IV CBASS anti-phage defense that involves direct conjugation between a protein N-terminus and a 7-deazaguanine nucleobase.

### Cap9 is a QueC-like enzyme adapted for protein modification

NDG is a direct nucleobase-protein conjugation that is chemically distinct from previously described protein modifications. To determine how type IV CBASS defense systems install NDG on the CD-NTase N-terminus, we purified CdnD from cells expressing *Rhizobiales* type IV CBASS operons with individual deletions of accessory genes encoding Cap9, Cap10, or Cap11. LC-MS analysis of purified CdnD revealed that deletion of Cap9 completely eliminated NDG conjugation, whereas deletion of Cap10 increased NDG conjugation, and deletion of Cap11 had no effect (Fig. 4A). Co-immunoprecipitation of CdnD from these mutant operons revealed that deletion of Cap9 reduced Cap10–CdnD complex formation, and deletion of Cap10 promoted binding of CdnD with a protein corresponding to the expected molecular weight of Cap9, which was confirmed by purification and LC-MS analysis of this protein complex (Fig. 4B, Fig. S7). Together these data suggest that NDG conjugation is catalyzed by Cap9 and that this modification is required to stabilize the Cap10–CdnD complex.

**Figure 4.**
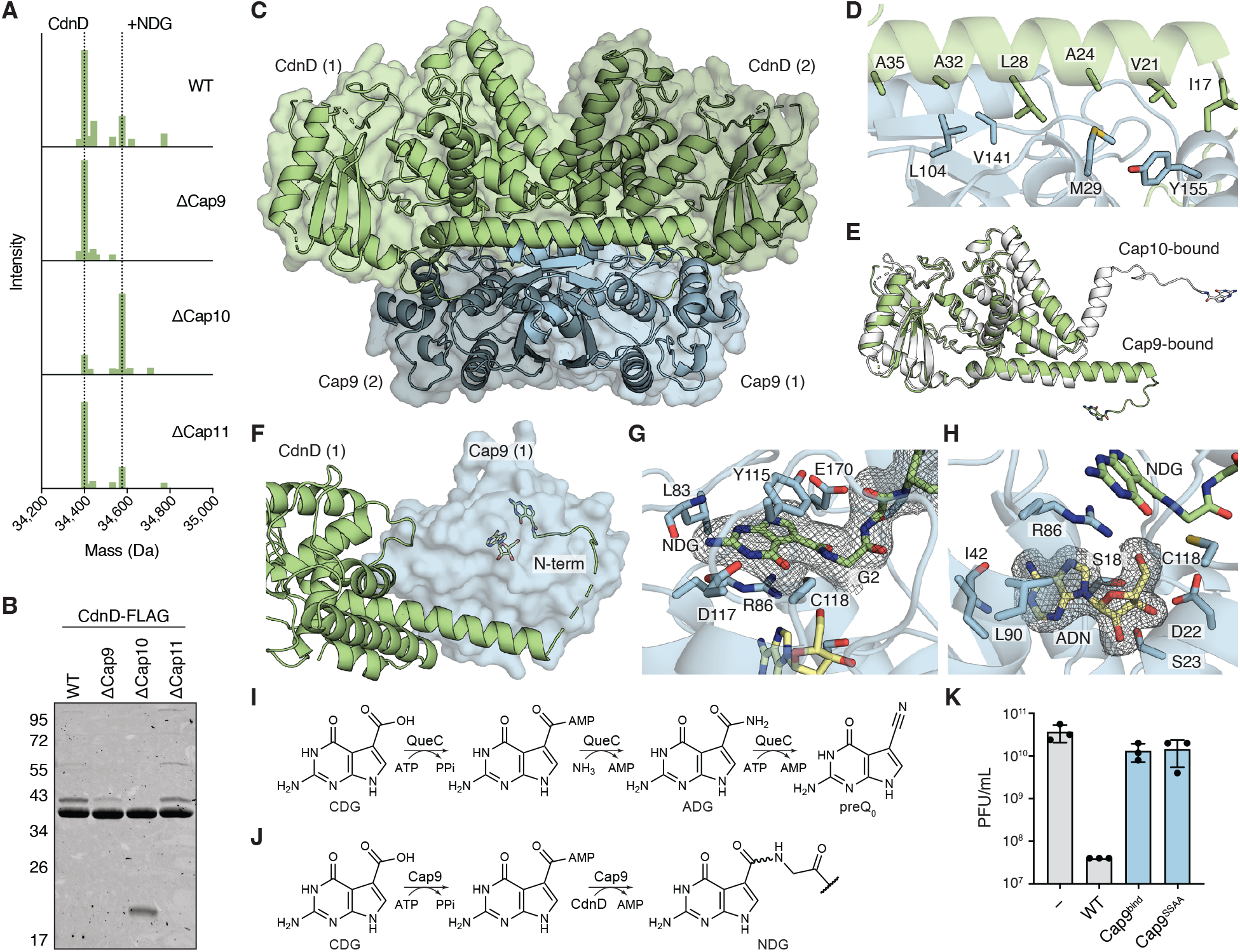
Cap9 catalyzes deazaguanylation through a QueC-like mechanism. (**A**) Intensity of ion masses determined by intact LC-MS analysis and spectral deconvolution of CdnD isolated from *E. coli* expressing *Rhizobiales* CBASS operons with the indicated genotypes. Dotted vertical lines show expected masses of methionine M1-cleaved CdnD with and without NDG modification. (**B**) Coomassie-stained SDS-PAGE following anti-FLAG immunoprecipitation. Lysates were prepared from *E. coli* expressing *Rhizobiales* operon encoding a C-terminal FLAG-tagged CdnD (CdnD-FLAG) and the indicated accessory protein deletions. Calculated molecular weights: CdnD-FLAG (34.5 kDa), Cap9 (19.9 kDa), Cap10 (37.4 kDa), Cap11 (23.5 kDa). (**C**) 2.2 Å X-ray crystal structure of *Rhizobiales* Cap9–CdnD complex. Two protomers of CdnD (green) and two protomers of Cap9 (blue) form a 2:2 heterotetrameric complex. Each Cap9–CdnD dimer is denoted with (1) or (2). See Table S2 for crystallographic statistics. (**D**) Hydrophobic interface between CdnD helix α1 (green) and Cap9 (blue). (**E**) Superposition of Cap10-bound CdnD structure (white) and Cap9-bound CdnD structure (green) reveals a large conformational change in the N-terminal tail and helix α1. (**F**) CdnD N-terminal tail (green) binds in Cap9 (blue) active site with adenosine (yellow). **G**) and **(H**) NDG-conjugated CdnD N-terminus (green) and adenosine (yellow) interact with residues inside the Cap9 active site (blue). Gray meshes depict F_O_−F_C_ polder omit maps (sigma=3). (**I**) Catalytic mechanism of QueC showing conversion of 7-carboxy-7-deazaguanine (CDG) to 7-amido-7-deazaguanine (ADG) and then to 7-cyano-7-deazaguanine (preQ_0_). (**J**) Proposed mechanism of Cap9-catalyzed NDG conjugation. CDG carboxylate is activated via reaction with ATP α-phosphate followed by nucleophilic attack with CdnD N-terminal amine to complete NDG conjugation. Bond connecting NDG and protein N-terminus indicated by wavy line. (**K**) Phage Bas18 plaque forming units (PFU) determined by plaque assay with *E. coli* expressing a GFP control (−) or *Rhizobiales* CBASS operon with the indicated genotypes. Wildtype (WT), Cap9 M29E/L104E/S135K (Cap9^bind^), Cap9 S18A/S23A (Cap9^SSAA^).

We determined a 2.2 Å X-ray crystal structure of CdnD isolated from cells lacking Cap10, revealing the structure of a Cap9–CdnD complex. Similar to the structure of Cap10–CdnD, the Cap9–CdnD complex is a symmetric 2:2 assembly with a core Cap9 homodimer flanked by two CdnD protomers (Fig. 4C, Table S2). Cap9 exhibits high overall structural similarity to the bacterial Q biosynthetic enzyme QueC, including a complete Rossman-like fold formed by a core of five parallel β-strands braced by five α-helices and a structural Zn^2+^-binding site (Fig. S8) (*36*). The active site of each CdnD protomer exhibits the same inactive conformation observed in the Cap10–CdnD structure with a rotated N-terminal lobe and misaligned metal-binding active site residues. In this structure, the coenzyme A binding site is occupied by a molecule of AMP (Fig. S8). The primary interface between Cap9 and CdnD occurs through a hydrophobic face of the CdnD helix α1 that docks into a complementary cleft along Cap9 (Fig. 4D). This helix α1 undergoes a large conformational change between the Cap9-bound state and Cap10-bound state where it also forms part of the major Cap10-binding interface (Fig. 4E). Mutations to Cap9 in the CdnD-binding cleft (M29E/L104E/S135K) abolish defense (Fig. 4K), highlighting the importance of Cap9–CdnD complex formation.

In the Cap9–CdnD assembly, the N-terminus of each CdnD protomer wraps around and inserts into the active site of Cap9 (Fig. 4F). Clear electron density allowed unambiguous assignment of both the NDG-conjugated CdnD N-terminus and an adjacent adenosine cofactor in the Cap9 active site that likely derives from AMP added during crystallization (Fig. 4G, 4H). The NDG modification is stabilized through hydrogen bonding with Cap9 residue D117 and the peptide backbone of L83, and aromatic stacking with Y115 and R86 (Fig. 4G). The adenosine molecule occupies an extended pocket below the NDG binding site, where the adenine nucleobase is coordinated through hydrogen bond interactions with the backbone of Cap9 residue I42, while residues S18, D22, and S23 form part of a conserved pyrophosphate-binding SXGXDS motif (Fig. 4H) (*37*).

Analysis of the Cap9–CdnD structure explains the mechanism of NDG conjugation. Canonical QueC proteins catalyze a two-step reaction to convert 7-carboxy-7-deazaguanine (CDG) to 7-amido-7-deazaguanine (ADG) and then to 7-cyano-7-deazaguanine (preQ_0_), consuming two molecules of ATP in the process (Fig. 4I) (*13, 38*). Notably, the Cap9–CdnD complex provides the first structure of a QueC-family enzyme bound to a 7-deazaguanine nucleobase and adenosine cofactor, revealing how the substrates are positioned to coordinate reaction between the nucleobase and adenosine α-phosphate or free ammonia. Ligand orientation within the structure explains how the Cap9 active site uses the same catalytic mechanism to direct nucleophilic attack by the CdnD N-terminal amine rather than ammonia into the AMP-activated carbonyl of CDG (Fig. 4J). Mutation of Cap9 pyrophosphate-binding residues S18 and S23 to alanine eliminated all anti-phage defense activity (Fig. 4K), explaining how a conserved reaction in Q biosynthesis is co-opted to enable nucleobase-protein conjugation in type IV CBASS immunity.

Type IV CBASS operons exhibit a striking evolutionary repurposing of 7-deazaguanine biosynthetic machinery from its canonical role in tRNA modification to nucleobase-protein conjugation in anti-phage immunity. In a series of structures, our results explain the function of Cap9 and Cap10 in CBASS anti-phage defense and reveal how these protein-modifying enzymes catalyze conjugation of an NDG modification that is chemically distinct from previously described post-translational modifications. In contrast to adenylation and ADP-ribosylation, where nucleotides are conjugated to proteins via connection through a phosphate or ribose (*39, 40*), NDG forms a direct linkage between the nucleobase and the protein. Additionally, unlike known nucleotide conjugations, NDG occurs at the protein N-terminus, a site of protein modification and regulation with unique chemical reactivity (*29, 41, 42*).

The functions of Cap9 and Cap10 in protein N-terminal deazaguanylation reveal an alternate reaction pathway that parallels ubiquitin-like conjugation systems. Compared to type II CBASS operons that rely on well characterized ubiquitin E1/E2-like enzymes to conjugate AMP to the CD-NTase C-terminus (*32*–*35*), the NDG nucleobase-protein conjugation in type IV CBASS achieves a similar regulatory mechanism using entirely distinct chemistry, targeting the CD-NTase N-terminus using Q biosynthetic pathway enzymes unrelated to ubiquitin ligase enzymes. The direct attachment of NDG to the CD-NTase N-terminus preserves the N9 position of the nucleobase, potentially allowing subsequent addition of ribose or integration into nucleic acid chains. This model provides a possible role for Cap11 as a negative regulator that reverses the NDG transfer to a target substrate, analogous to the mechanism by which the predicted Cap11 homolog OGG excises oxidized guanine bases from DNA (*43*). Although the final target substrate in type IV CBASS remains unknown, the discovery of NDG now allows comparative analyses of type II CBASS and type IV CBASS operons to illuminate this critical step in CD-NTase activation across bacterial defense systems. Finally, the incredibly widespread distribution of QueC-like proteins throughout bacteria and archaea suggests that there may be numerous additional systems where enzymes like Cap9 have been repurposed from nucleobase modification to protein targeting.

## MATERIALS AND METHODS

### Bacterial strains and phages

*E. coli* strain BW25113 was grown in lysogeny broth (LB) supplemented with 5 mM MgCl_2_, 5 mM CaCl_2_, 0.1 mM MnCl_2_, and 100 µg mL^−1^ ampicillin with 1.5% agar for bacterial plates, 0.5% low-melt agar for top agar, or no agar for liquid cultures. Phages were propagated by picking a single phage plaque into a liquid bacterial culture grown to optical density at 600 nm (OD_600_) of 0.3 and incubating with shaking at 37°C until culture collapse. The culture was then centrifuged for 10 min at 3,200 × g, and the supernatant was filtered through a 0.2 µm filter and stored at 4°C. The titer of the lysate was determined using the plaque assay described below.

### Plasmid construction

Native type IV CBASS operon sequences were synthesized and cloned into an arabinose-inducible pBAD vector containing an ampicillin resistance cassette by GenScript. Mutations, deletions, and C-terminal FLAG or 6×His tags were introduced using site-directed mutagenesis PCR and sequences were confirmed by whole plasmid sequencing. Gene deletions were designed to truncate the majority of the coding sequence, leaving ten N-terminal residues and ten C-terminal residues to prevent disruption of translation of overlapping neighboring coding sequences.

### Phage plaque assay

Liquid cultures were inoculated from single bacterial colonies or glycerol stocks and incubated with shaking at 37°C for 6 h until OD_600_ reached at least 0.3. Cultures were diluted to OD_600_=0.06 in melted top agar supplemented with L-arabinose and poured onto a bacterial plate supplemented with L-arabinose (6 mL top agar for 9 cm round plate, 15 mL top agar for 12 cm square plate) and allowed to solidify at room temperature for 1 h. L-arabinose was supplemented at 0.2% (w/v) for initial screening and then reduced to 0.002% (w/v) for subsequent experiments to reduce operon toxicity. Phages were prepared in 10-fold serial dilutions from high titer stocks with SM buffer (50 mM Tris pH 7.5, 100 mM NaCl, 8 mM MgSO_4_) and spotted onto the cooled top agar in 2.5 µL drops. Plates were incubated at room temperature for 30 min to dry and incubated overnight at 30°C. Plates were imaged using a Bio-Rad ChemiDoc MP and phage plaques were counted manually from images. In cases where individual phage plaques could not be distinguished, the highest dilution phage drop with observable bacterial clearance was counted as 10 plaques.

### Bacterial toxicity

Liquid cultures were inoculated from single bacterial colonies and incubated with shaking at 37°C for 6 h until OD_600_ reached at least 0.3. Bacteria were pelleted by centrifugation and resuspended at OD_600_=0.2 in phosphate-buffered saline (PBS). 10-fold serial dilutions were prepared with PBS, and diluted bacteria were spotted in 2.5 µL drops onto plates supplemented with L-arabinose or D-glucose. Plates were dried at room temperature for 30 min and incubated overnight at 30°C. Plates were imaged using a Bio-Rad ChemiDoc MP.

### FLAG immunoprecipitation

Liquid cultures (20 mL) were inoculated from single bacterial colonies and incubated with shaking at 37°C until OD_600_ reached 1. L-arabinose was added to a final concentration of 0.2% (w/v) and the culture was incubated overnight with shaking at 37°C. After overnight expression, cell pellets were collected by centrifugation, resuspended in 1 mL cold TBS (50 mM Tris pH 7.5, 150 mM NaCl) and lysed by sonication. Lysate was cleared by centrifugation at 20,000 × g for 30 min. Anti-FLAG M2 Magnetic Beads (Millipore Sigma) were washed and resuspended in TBS, and 20 µL packed beads were added to each 1 mL cleared lysate sample. Samples were incubated with gentle rolling for 2 h at 4°C and beads were then washed three times with TBS and eluted with 50 µL of 150 µg mL^−1^ 3×FLAG peptide (APExBIO) in TBS.

### Recombinant protein expression and purification

Overnight liquid cultures of bacteria were inoculated from single bacterial colonies transformed with plasmid containing the *Rhiziobiales* CBASS operon encoding a C-terminal 6×His tagged CdnD. Liquid cultures (2 × 1L) of 2YT (16 g L^−1^ Bacto tryptone, 10 g L^−1^ yeast extract, 5 g L^−1^ NaCl, pH 7.0, 100 µg mL^−1^ ampicillin) were inoculated with 1 mL of overnight culture and incubated with shaking at 37°C until OD_600_ reached 1. L-arabinose was added to a final concentration of 0.2% (w/v) and cultures were cooled to 16°C and incubated overnight with shaking. After overnight expression, cell pellets were collected by centrifugation and then resuspended and lysed by sonication in 60 mL lysis buffer (20 mM HEPES-KOH pH 7.5, 400 mM NaCl, 10% glycerol, 30 mM imidazole). Lysate was cleared by centrifugation at 40,000 × g for 30 minutes, supernatant was poured over 8 mL Ni-NTA resin (Qiagen), resin was washed twice with 35 mL lysis buffer, and protein was eluted with elution buffer (20 mM HEPES-KOH pH 7.5, 400 mM NaCl, 10% glycerol, 250 mM imidazole). Samples were then purified by size-exclusion chromatography using a 16/600 Superdex 200 column (Cytiva) using gel filtration buffer (20 mM HEPES-KOH pH 7.5, 250 mM KCl). For samples not used for crystallography, gel filtration buffer was supplemented with 10% glycerol. Purified proteins were concentrated to 10 mg mL^−1^ using 10-kDa MWCO centrifugal filter units (Millipore Sigma), aliquoted, flash frozen in liquid nitrogen, and stored at −80°C. For proteins purified under reducing conditions 1 mM TCEP was supplemented into all buffers.

### X-ray crystallography

Recombinant Cap10–CdnD complex was expressed and purified from cells containing *Rhiziobiales* CBASS operon using a C-terminal 6×His CdnD tag, and Cap9–CdnD complex was purified similarly using a ΔCap10 mutant operon. Crystals were grown in hanging-drop format at 18°C. Crystals of Cap10–CdnD were grown in 2 µL drops containing a 1:1 mixture of 5 mg mL^−1^ protein in buffer containing 20 mM HEPES-KOH pH 7.5 and 75 mM KCl and reservoir solution containing 0.1 M HEPES-KOH pH 7.5, 0.3 M calcium acetate, and 10% PEG-8000 and cryo-protected with reservoir solution supplemented with 30% ethylene glycol. Crystals of Cap9–CdnD were grown in 2 µL drops containing a 1:1 mixture of 5 mg mL^−1^ protein in buffer containing 20 mM HEPES-KOH pH 7.5, 75 mM KCl, 5 mM MgCl_2_, and 1 mM AMP and reservoir solution containing 0.1 M Tris-HCl pH 8.5 and 24% PEG-3350 and cryo-protected with reservoir solution supplemented with 16% ethylene glycol, 5 mM MgCl_2_, and 1 mM AMP. X-ray diffraction data were collected at National Synchrotron Light Source II beamline 17-ID-2 and processed using autoPROC (*44*). Initial search models were predicted using AlphaFold2 (*45*), molecular replacement and model refinement were performed using PHENIX (*46*), and model building was performed using Coot (*47*). Crystallographic statistics are provided in Table S2 and all structural figures were generated using PyMOL (Schrödinger).

### Mass spectrometric data collection and spectral deconvolution of intact proteins

Recombinant CdnD alone or in complex with Cap9 or Cap10 was expressed and purified using a C-terminal 6×His CdnD tag as described above. Normalized protein samples were diluted to 1 µM in CH3CN/H2O (1:1) and analyzed by intact protein LC/MS using a Waters Xevo G2-XS QTof system equipped with an Acquity UPLC BEH Premier C4 1.7 µm column. The mobile phase was a linear gradient of 5–95% acetonitrile / water + 0.05% formic acid. Peak integration and spectral deconvolution (MaxEnt1, iterated to convergence at 1 Da resolution) was performed using MassLynx and OpenLynx software packages.

### Mass spectrometric data collection and analysis of CdnD N-terminal peptide

Recombinant Cap10–CdnD complex was subjected to SDS-PAGE and the band corresponding to CdnD was cut from the gel, resuspended in 200 mM EPPS pH 8.5 and digested with trypsin at 100:1 protein-to-protease ratio for 6 h at 37°C. The sample was desalted via StageTip, dried again via vacuum centrifugation, and reconstituted in 5% acetonitrile, 5% formic acid for LC-MS/MS processing (*48*). Mass spectrometry data were collected using an Orbitrap Astral mass spectrometer (Thermo Fisher Scientific, San Jose, CA) coupled with a Neo Vanquish liquid chromatograph. Orbitrap detector of MS2 was used. Peptides were separated on a 110 cm uPAC C18 column (Thermo Fisher Scientific). For each analysis, ∼0.5 µg protein was loaded onto the column. Peptides were separated using a 60 min gradient of 7–24% acetonitrile in 0.125% formic acid with a flow rate of 400 nL min^−1^. The scan sequence began with an MS1 spectrum (Orbitrap analysis, resolution 60,000, 350–1360 Th, automatic gain control (AGC) target set to 500%, maximum injection time set to 50 ms). The hrMS2 stage consisted of fragmentation by higher energy collisional dissociation (HCD, normalized collision energy 27%) and analysis using the Astral mass analyzer (AGC 200%, maximum injection time 15 ms, isolation window 1.2 Th). Data were acquired using the FAIMSpro interface with the dispersion voltage (DV) set to 5,000 V, the compensation voltages (CVs) set to −30 V, −40 V, −50 V, and −60 V or −40 V, −60V, and −70V. The TopSpeed parameter was set at 1 sec per CV. Mass spectra were processed using a Comet-based in-house software pipeline. MS spectra were converted to mzXML using a modified version of ReAdW.exe. Database searching included all entries from the human UniProt database, which was concatenated with a reverse database composed of all protein sequences in reversed order. Searches were performed using a 3 Da precursor ion tolerance. Product ion tolerance was set to 0.03 Th, while considering the following parameters: XCorr > 1.5 and a mass shift of 176 Th.

### In vitro CD-NTase activity

Reactions of 100 µL containing 0.5 mM each ATP, GTP, CTP, UTP with 10 mM MgCl_2_, 1 mM MnCl_2_, 20 mM Tris-HCl pH 7.5, 100 mM KCl, and 5 mM recombinant protein or buffer control (20 mM HEPES-KOH, 250 mM KCl, 10% glycerol) were incubated at 37°C for 4 h. Reactions were quenched by addition of 5 units of Quick CIP (New England Biolabs) and incubation at 37°C for 30 min followed by filtration through a 10-kDa MWCO spin filter (Millipore Sigma). Reactions were analyzed using an Agilent 1260 HPLC equipped with a diode array detector and an Agilent 6125 single quadrupole mass spectrometer in negative mode using a reverse-phase Agilent InfinitiLab Poroshell SB-Aq column (2.7 µm particle size, 2.1 mm inner diameter, 100 mm length) at a flow rate of 0.45 mL min^−1^ with a gradient from 100% solvent A (0.1% (w/v) ammonium formate) to 100% solvent B (methanol).

### Alignment of CD-NTase sequences

Protein sequences were aligned using MAFFT with default L-INS-I parameters (*49*). Sequence logo diagrams were generated with WebLogo 3 (*50*) using the initial eight residues of type IV CBASS CD-NTase sequences. Sequence alignments were visualized with Jalview (*51*).

## ACKNOWLEDGEMENTS

The authors are grateful to members of the Kranzusch laboratory for helpful comments and discussion. We thank D. Richmond-Buccola from the Kranzusch laboratory for assistance with phage challenge assays, and S. Mooney, H. Toyoda, and A. Lu from the Kranzusch laboratory for assistance with protein purification and crystallization. The work was funded by grants to P.J.K. from the Pew Biomedical Scholars program, the Burroughs Wellcome Fund PATH program, The G. Harold and Leila Y. Mathers Charitable Foundation, The Mark Foundation for Cancer Research, the Cancer Research Institute, the Parker Institute for Cancer Immunotherapy, and the National Institutes of Health (1DP2GM14650-01). K.M.S. thanks a gift from the Sjöberg Foundation. This work was funded in part by grants from the National Institutes of Health to J.A.P. (GM132129) and S.P.G. (GM97645). D.R.W. is supported through a Helen Hay Whitney Foundation postdoctoral fellowship. P.P. was supported by a postdoctoral fellowship from the Swiss National Science Foundation (SNF Mobility Grant P500PN_214278). X-ray data were collected at The Center for Bio-Molecular Structure (CBMS) that is primarily supported by the NIH-NIGMS through a Center Core P30 Grant (P30GM133893), and by the DOE Office of Biological and Environmental Research (KP1607011). NSLS2 is a U.S. DOE Office of Science User Facility operated under Contract No. DE-SC0012704. This publication resulted from the data collected using the beamtime obtained through NECAT BAG proposal #311950.

## AUTHOR CONTRIBUTIONS

The study was designed and conceived by D.R.W. and P.J.K. Phage challenge assays, biochemical experiments, and crystallography experiments and structure modeling were performed by D.R.W. Intact protein mass-spectrometry was performed by P.P. and K.M.S. NDG LC-MS/MS analysis was performed by J.A.P. and S.P.G. The manuscript was written by D.R.W. and P.J.K. All authors contributed to editing the manuscript and support the conclusions.

## DATA AND MATERIALS AVAILABILITY

Coordinates and structure factors of the Cap10–CD-NTase and Cap9–CD-NTase complexes have been deposited in PDB under the accession codes 9NTN, and 9NTO. All other data are available in the main text or the supplementary information. Correspondence and requests for materials should be addressed to P.J.K.

## SUPPLEMENTARY MATERIALS

Supplementary Figures S1 to S8

Supplementary Tables S1 and S2

