## Supplemental Figures for "Deazaguanylation is a nucleobase-protein conjugation required for type IV CBASS immunity"

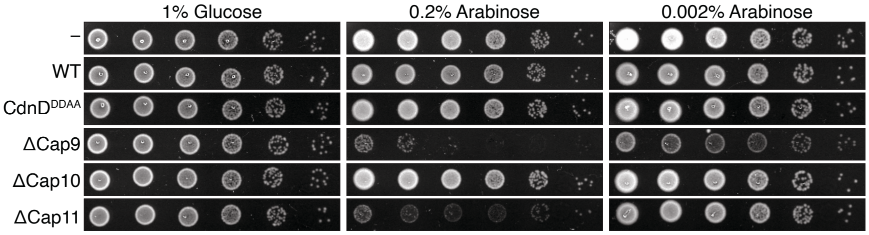


**Supplemental Figure S1 | Toxicity of mutant type IV operons.** Spot assay of *E. coli* containing a GFP control (−) or *Rhizobiales* CBASS operon with the indicated genotypes. Expression is repressed with 1% glucose or induced with 0.2% or 0.002% arabinose.

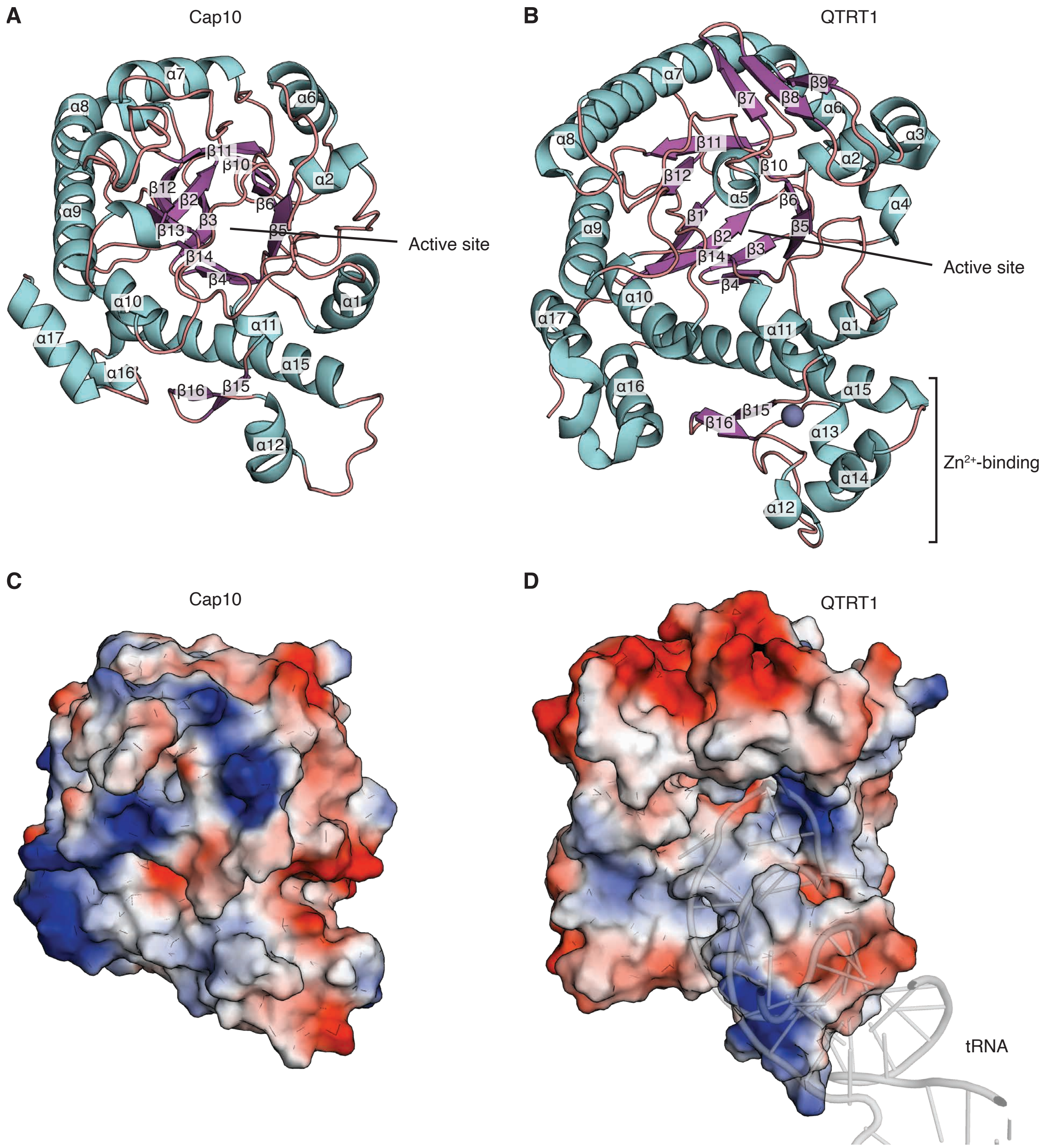


**Supplemental Figure S2 | Structural analysis of Cap10.** (**A**) Annotated secondary structure of Cap10. (**B**) Annotated secondary structure of QTRT1. PDB: 8OMR. (**C**) Calculated surface electrostatic potential of Cap10. (**D**) Calculated surface electrostatic potential of QTRT1 with bound tRNA shown.

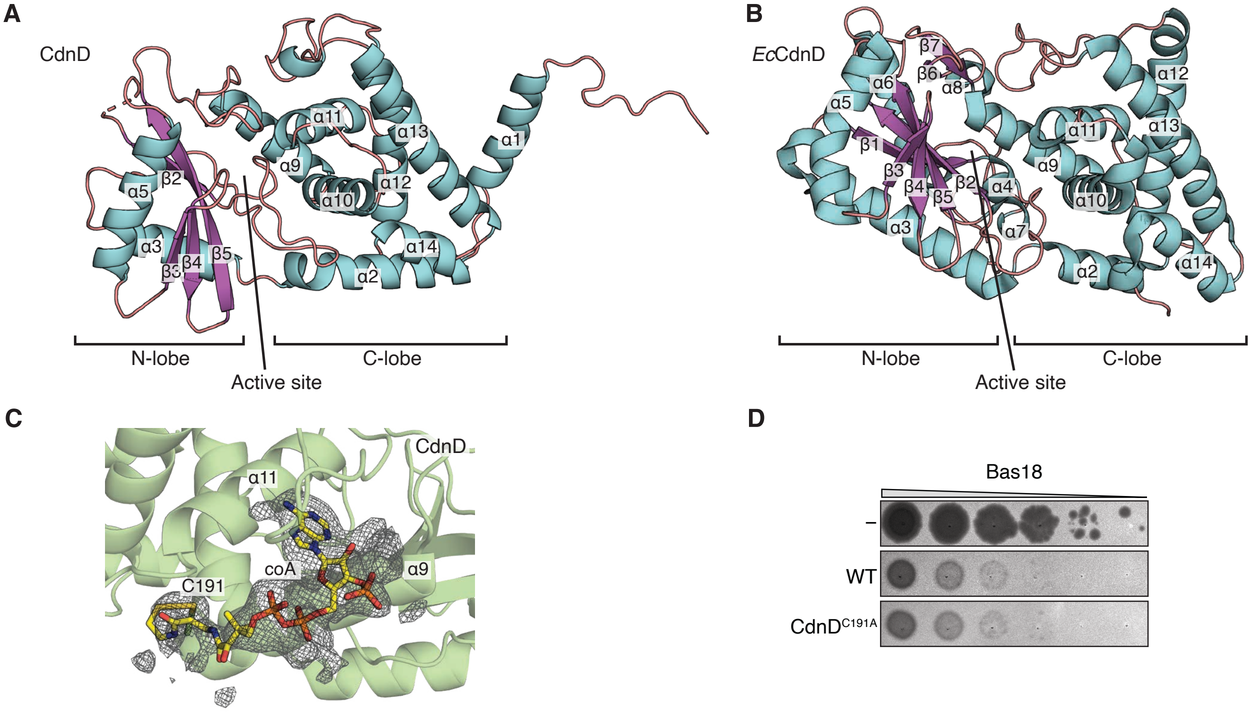


**Supplemental Figure S3 | Structural analysis of CdnD.** (**A**) Annotated secondary structure of CdnD from *Rhizobiales* Cap10–CdnD complex. (**B**) Annotated secondary structure of *Enterobacter cloacae* CdnD. PDB: 7LJL. (**C**) Coenzyme A ligand bound to CdnD, forming a disulfide bridge with residue C191. Gray mesh depicts $F_O-F_C$ polder omit map (sigma=3). (**D**) Representative plaque assay using phage Bas18 with *E. coli* expressing a GFP control (−) or *Rhizobiales* CBASS operon with the indicated genotype.

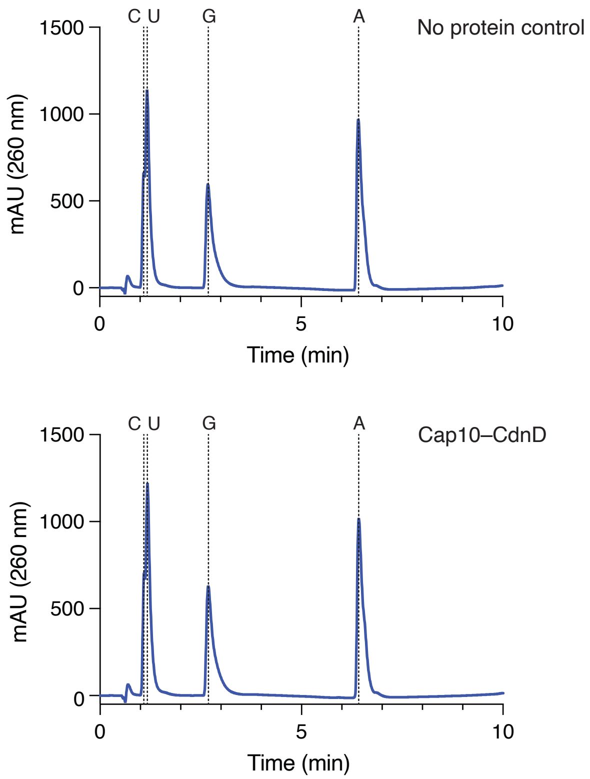


**Supplemental Figure S4 | Cap10–CdnD complex does not catalyze cyclic nucleotide synthesis.** HPLC trace showing UV absorbance at 260 nm of *in vitro* reactions containing Cap10–CdnD protein complex or buffer control. Reactions were quenched with calf-intestine phosphatase (CIP) before HPLC analysis. Elution time of uridine (U), cytidine (C), guanosine (G), and adenosine (A) were determined using chemical standards and mass spectrometry.

**A**

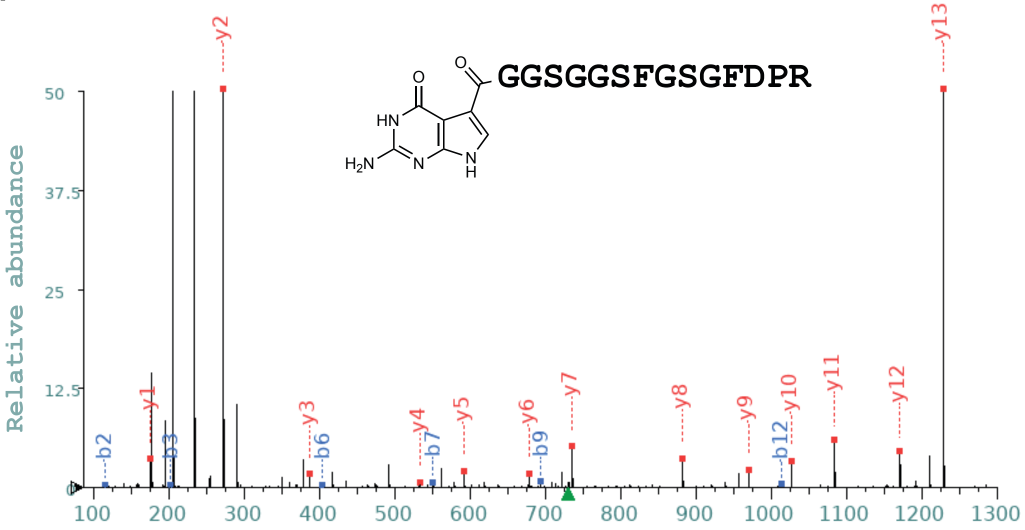


**B**

| Seq | # | b: Δ Error | b | y | y: Δ Error | +1 |
|---|---|---|---|---|---|---|
| G | 1 | --- | 58.029 | --- | --- | 14 |
| G | 2 | 38.575 | 115.050 | 1227.539 | 1.134 | 13 |
| S | 3 | 18.337 | 202.082 | 1170.517 | 1.797 | 12 |
| G | 4 | --- | 259.104 | 1083.485 | 1.196 | 11 |
| G | 5 | --- | 316.125 | 1026.464 | 0.409 | 10 |
| S | 6 | 12.435 | 403.157 | 969.442 | 1.608 | 9 |
| F | 7 | 0.859 | 550.226 | 882.410 | 0.920 | 8 |
| G | 8 | --- | 607.247 | 735.342 | 0.182 | 7 |
| S | 9 | 4.161 | 694.279 | 678.321 | 1.067 | 6 |
| G | 10 | --- | 751.301 | 591.289 | 1.611 | 5 |
| F | 11 | --- | 898.369 | 534.267 | 0.602 | 4 |
| D | 12 | 0.442 | 1013.396 | 387.199 | 0.578 | 3 |
| P | 13 | --- | 1110.449 | 272.172 | 0.583 | 2 |
| R | 14 | --- | --- | 175.119 | 0.900 | 1 |

**Supplemental Figure S5 | LC-MS/MS fragmentation analysis of modified CdnD N-terminal peptide.** (**A**) Extracted MS/MS spectrum of modified CdnD N-terminal peptide. (**B**) Table of fragment ions and associated mass errors.

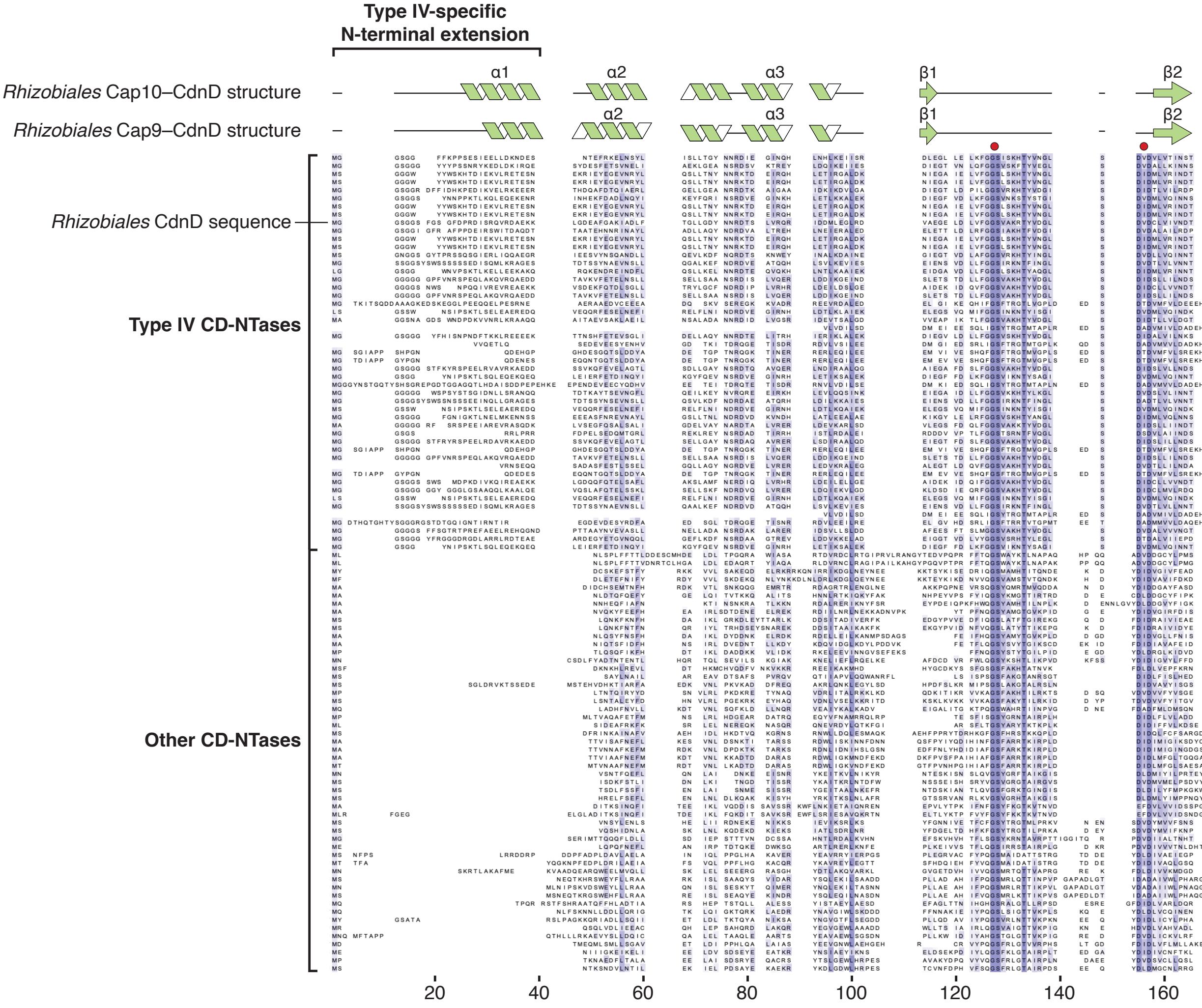


**Supplemental Figure S6 | Sequence alignment of CD-NTase proteins.** Sequence alignment of N-terminal segments from type IV CBASS CD-NTases and other representative CD-NTases (*20*). Secondary structure is annotated from *Rhizobiales* CdnD structure in complex with Cap10 or Cap9. Red dots denote conserved GS and DXD active site motifs. Amino acid conservations depicted with blue shading.

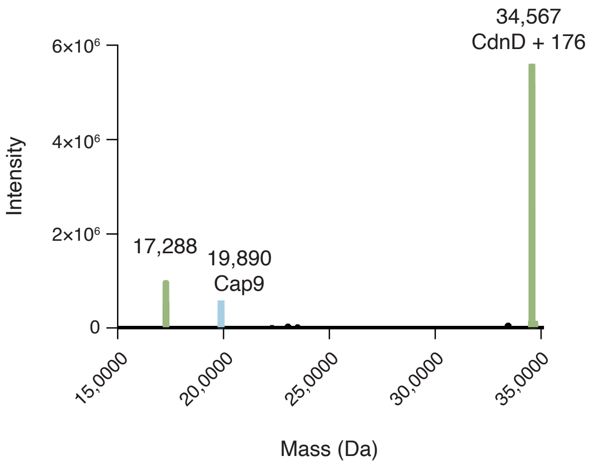


**Supplemental Figure S7 | LC-MS analysis of Cap9–CdnD complex.** Intensity of ion masses determined by intact LC-MS analysis and spectral deconvolution of Cap9–CdnD complex isolated from *E. coli* expressing ΔCap10 mutant *Rhizobiales* CBASS operon. Parent and half-mass peaks for NDG-modified CdnD shown in green (34,567 and 17,284 Da) and peaks for Cap9 mass (19,893 Da) shown in blue.

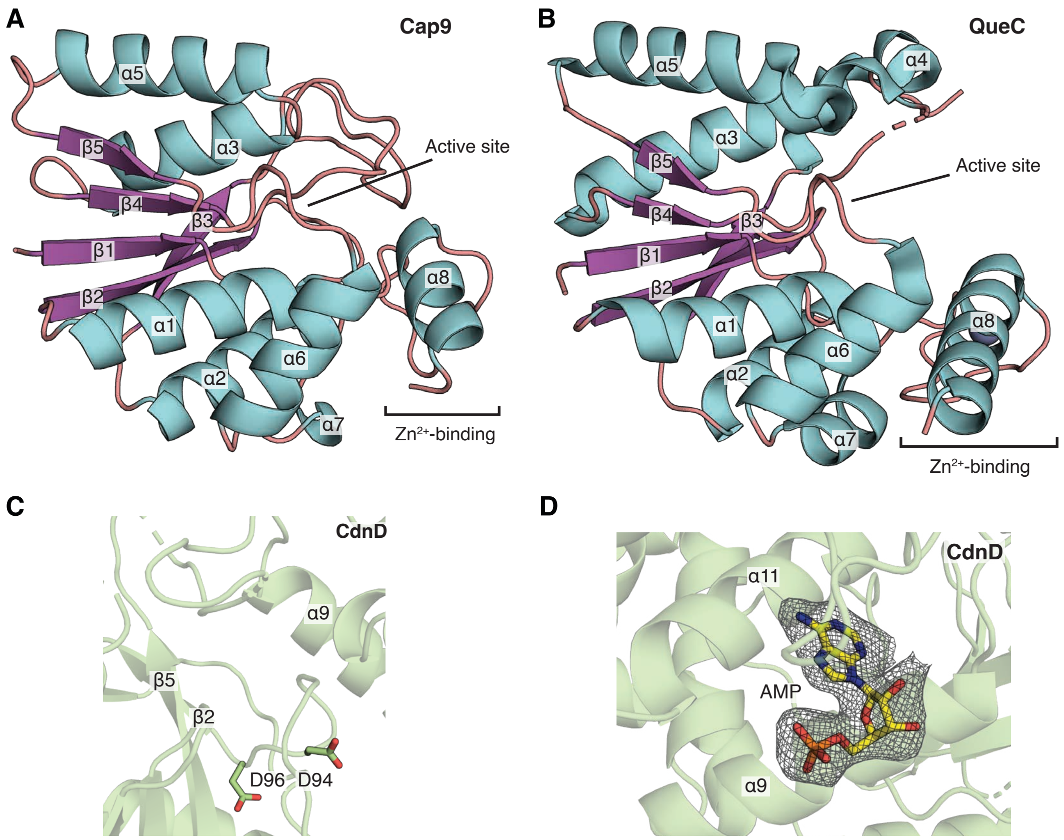


**Supplemental Figure S8 | Structural analysis of Cap9–CdnD complex.** (**A**) Annotated secondary structure of Cap9. (**B**) Annotated secondary structure of *Bacillus subtilis* QueC. PDB: 3BL5. (**C**) CdnD active site with metal-binding residues D94 and D96. (**D**) AMP (yellow) bound to CdnD in Cap9–CdnD complex in identical binding site as coenzyme A in Cap10–CdnD structure. Gray mesh depicts $F_O - F_C$ polder omit map (sigma=3).
